# Quantifying the fitness benefit and cost of cefixime resistance in *Neisseria gonorrhoeae* to inform prescription policy

**DOI:** 10.1101/121418

**Authors:** Lilith K Whittles, Peter J White, Xavier Didelot

## Abstract

Gonorrhea is one of the most common bacterial sexually transmitted infections in England. Over 41,000 cases were recorded in 2015, more than half of which occurred in men who have sex with men (MSM). As the bacterium has developed resistance to each first-line antibiotic in turn, we need an improved understanding of fitness benefits and costs of antibiotic resistance to inform control policy and planning. Cefixime was recommended as a single dose treatment for gonorrhea from 2005 to 2010, during which time resistance increased and subsequently declined. We developed a stochastic compartmental model representing the natural history and transmission of cefixime sensitive and resistant strains of *Neisseria gonorrhoeae* in MSM in England, which was applied to data on diagnoses and prescriptions between 2008 and 2015. We estimated that asymptomatic carriers play a crucial role in overall transmission dynamics, with about 40% of infections remaining asymptomatic and untreated, accounting for 96% of onward transmission. The fitness cost of cefixime resistance in the absence of cefixime usage was estimated to be such that the number of secondary infections caused by resistant strains is only about half as much as for the susceptible strains, which is insufficient to maintain persistence. However, we estimated that treatment of cefixime-resistant strains with cefixime was unsuccessful in 84% of cases, representing a fitness benefit of resistance. This benefit was large enough to counterbalance the fitness cost when 31% of cases are treated with cefixime, and when more than 51% of cases were treated with cefixime the resistant strain had a net fitness advantage over the susceptible strain. Our findings have important implications for antibiotic stewardship and public health policies, and in particular suggest that cefixime could be used to treat a minority of gonorrhea cases without raising resistance levels.

## Introduction

Gonorrhea, caused by the bacterial pathogen *Neisseria gonorrhoeae*, is one of the most common sexually transmitted infections in England. Incidence has increased year on year since 2008, culminating in over 41,000 cases in 2015 [1]. Around 22,000 of these cases were found in men who have sex with men (MSM), constituting a 20% annual increase. The greatest cause for concern, however, is the rapid growth in antimicrobial resistance. The bacterium has quickly developed resistance to each first-line antibiotic in turn, from penicillin through to cephalosporins, such as cefixime and ceftriaxone [2]. Treatment with ceftriaxone is the last remaining single-drug option in most settings worldwide; however susceptibility is diminishing rapidly [3]. As a result, England and many other countries now recommend treatment of gonorrhea with a dual therapy of ceftriaxone and azithromycin [4]. Ceftriaxone resistance has been detected only sporadically in England; however azithromycin resistance is easily selected for and was prevalent in a recent outbreak [5]. Resistance to azithromycin effectively reduces the current treatment to a monotherapy, making resistance trends increasingly important to monitor against the threat of potentially untreatable gonorrhea.

Public Heath England (PHE) runs the Gonococcal Resistance to Antimicrobials Surveillance Programme (GRASP) [68], which has produced a report annually since 2000 [6–21]. GRASP monitors trends in resistance and susceptibility to a panel of antibiotics used to treat gonorrhea in England and Wales, and thus informs national treatment guidelines and strategy. In 2004, GRASP began testing for cefixime resistance, defined as having a Minimum Inhibitory Concentration (MIC) of ≥ 0.125 mg/l [10]. In 2005, following worrying increases in resistance to the previous therapy, ciprofloxacin, a new recommendation was introduced that uncomplicated gonorrhea should be treated with a single dose of cefixime [22].

Fig 1 shows the trends in cefixime prescription and resistance in England. Very little resistance was detected until 2007, however by 2009 the total level of resistance had passed the 5% threshold at which the WHO recommends that first-line treatment guidelines should be changed [13,23]. At this time almost 60% of gonorrhea diagnoses were being treated with cefixime [15]. The majority of the resistance was concentrated in the MSM population, where it reached a peak of 33% in 2010 [16]. This evidence, combined with increasingly common reports of cefixime treatment failure, formed the basis for the decision in May 2011 for another update to the treatment guidelines for uncomplicated gonorrhea [24,25]. Cefixime was no longer recommended as a first-line treatment, and was replaced with a combination of 500mg ceftriaxone and 1g azithromycin [4]. Since 2011, cefixime prescribing has fallen drastically, in line with the updated guidelines. Over the same period the proportion of cefixime-resistant isolates has declined steadily in MSM, falling to less than 1% in 2014 [20].

**Fig 1.**
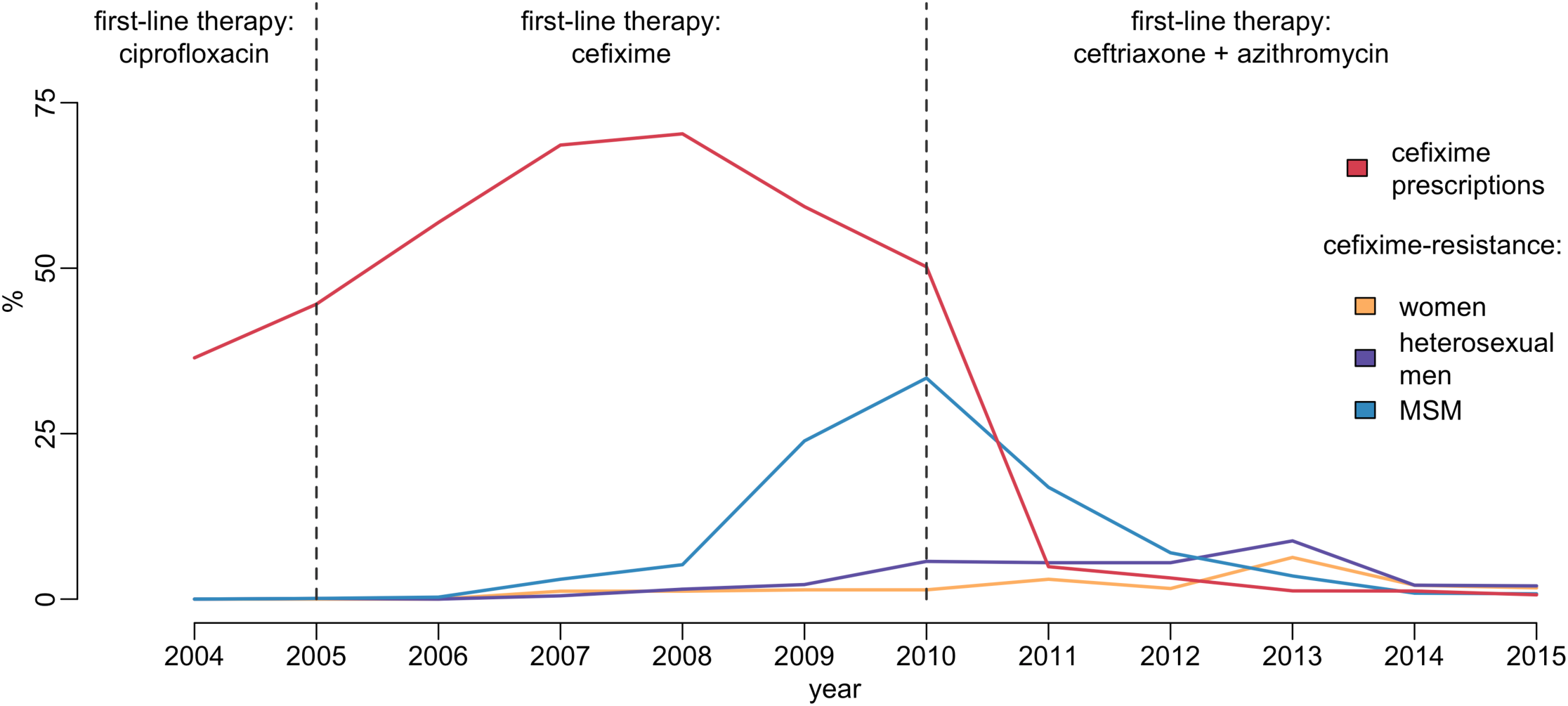
Usage and resistance of cefixime in England and Wales. The proportion of gonococcal isolates in GRASP resistant to cefixime over time is compared with the proportion of gonorrhea diagnoses treated with cefixime. Dashed lines show dates of treatment guideline changes.

We hypothesize that the resistance trend observed can be explained by a net fitness benefit to cefixime resistance when cefixime is widely prescribed but a net fitness cost when cefixime prescriptions decline. Understanding the relationship between antibiotic use and the emergence of resistance in gonorrhea has been identified as a key research agenda [26]. Here our main aim is to further the understanding of the evolutionary dynamics of cefixime resistance, and to use this newfound knowledge to inform public health practice. There is still much we do not understand about the natural history of gonorrhea, especially since unobserved asymptomatic infections have long been thought to be an important reservoir of infection. The proportion of incident cases that are asymptomatic at each bodily site of infection is known to vary, but has not been definitively measured [27–29]. Furthermore, the expected duration of carriage of asymptomatic gonococcal infection is not well studied. Estimates have been traditionally in the region of six months, however recent work using genomic data on pairs of known sexual contacts has suggested that a longer duration of carriage can occasionally happen [30]. We therefore developed and applied a Bayesian statistical approach to account for these uncertainties in the epidemiology of gonorrhea.

## Materials and methods

### Epidemiological Data

The total number of diagnoses of gonorrhea in MSM in England between 2008 and 2015 were extracted from the Genitourinary Medicine Clinic Activity Dataset (GUMCAD) [60]. This mandatory reporting system provides data on diagnoses of sexually transmitted infections from sexual health services in England, and the GUMCAD data is published annually by PHE on their website at http://www.gov.uk/government/statistics/sexually-transmitted-infections-stis-annual-data-tables. This yearly number of gonorrhea diagnoses is denoted *Y*(*t*).

The number of cases of gonorrhea in MSM that were cefixime-resistant and reported by GRASP between 2008 and 2015 were extracted from the corresponding GRASP reports [14–21] and denoted *Y*^res^(*t*). The coverage of GRASP was calculated for every year between 2008 and 2015 by taking the ratio between the number of cases included in GRASP (irrespective of resistance) and the number of GUMCAD diagnoses in the same year. This GRASP coverage proportion is denoted *q*(*t*). The proportion of gonorrhea cases that were treated with cefixime, as opposed to other antibiotics, was also extracted from the GRASP reports between 2008 and 2015. This time-dependent proportion is denoted π(*t*) and illustrated in Fig 1.

The MSM population is estimated at *N* = 1.5 million, based on a sexually active male population of 20 million [33]. In the third National Survey of Sexual Attitudes and Lifestyles (Natsal) 8.4% of men reported same-sex experience at least once, with 2.6% of men having had a same-sex partner in last five years, putting a plausible range for the MSM population at 0.5 and 1.7 million [34]. Natsal is known to under-represent MSM so an estimate towards the top of the range was adopted [35]. Given the low prevalence of gonococcal infection in the population, the total population size is not expected to excessively affect the results.

### Transmission model

In order to investigate the fitness cost and benefit of cefixime resistance in gonorrhea, we created a stochastic compartmental model, illustrated in Fig 2 with notation summarised in Table 1. High rates of reinfection with gonorrhea have been observed, suggesting low levels of acquired immunity [31], and experimental urethral infection in male volunteers found no protection was conferred on repeat infection with an identical strain six months apart [32]. It was therefore assumed that no immunity was conferred upon recovery from infection. The analysis was restricted to MSM, the population in which the cefixime-resistant outbreak of gonorrhea was concentrated. A closed population of size *N* was assumed due to the short time period under consideration. Individuals are initially susceptible (*S*). They become infected with strain *s* ϵ{sus, res}, denoting cefixime-susceptible and resistant strains respectively. The model assumes that strains do not vary in transmissibility, and that the rate of infection from an infectious individual to a susceptible individual is *θ*/*N*. Infected individuals initially pass through an incubation period (*U_s_*) which they leave at rate *σ*. A proportion *ψ* of those infected then go on to develop symptoms (*E_s_*), whereas the remainder enters an asymptomatic stage (*A_s_*). Recovery from asymptomatic infection happens (either naturally or following unrelated antibiotic treatment) at rate *v* for the susceptible strain and at rate *αv* for the resistant strain. The parameter α therefore represents the fitness cost of cefixime resistance. The infected population for each strain *s* is denoted *I_s_* = *U_s_* + *E_s_* + *A_s_*, and the total infected population denoted *I* = *I*_sus_ + *I*_res_. All infected individuals are assumed to be infectious. The symptomatic individuals (*E_s_*) seek treatment at rate *μ*. A time-varying proportion *π*(*t*) are treated with cefixime (*T*_s;cef_) whereas the remaining 1 – *π*(*t*) are treated with other antibiotics (*T*_s;oth_). The treated individuals recover from the infection and become susceptible again at rate *ρ*, with the exception of a proportion *ϕ* of the individuals infected with a cefixime-resistant strain and who have been treated with cefixime (*T*_res;cef_) for whom treatment failure happens and who become asymptomatically infected (*A*_res_) [51, 69].

**Fig 2.**
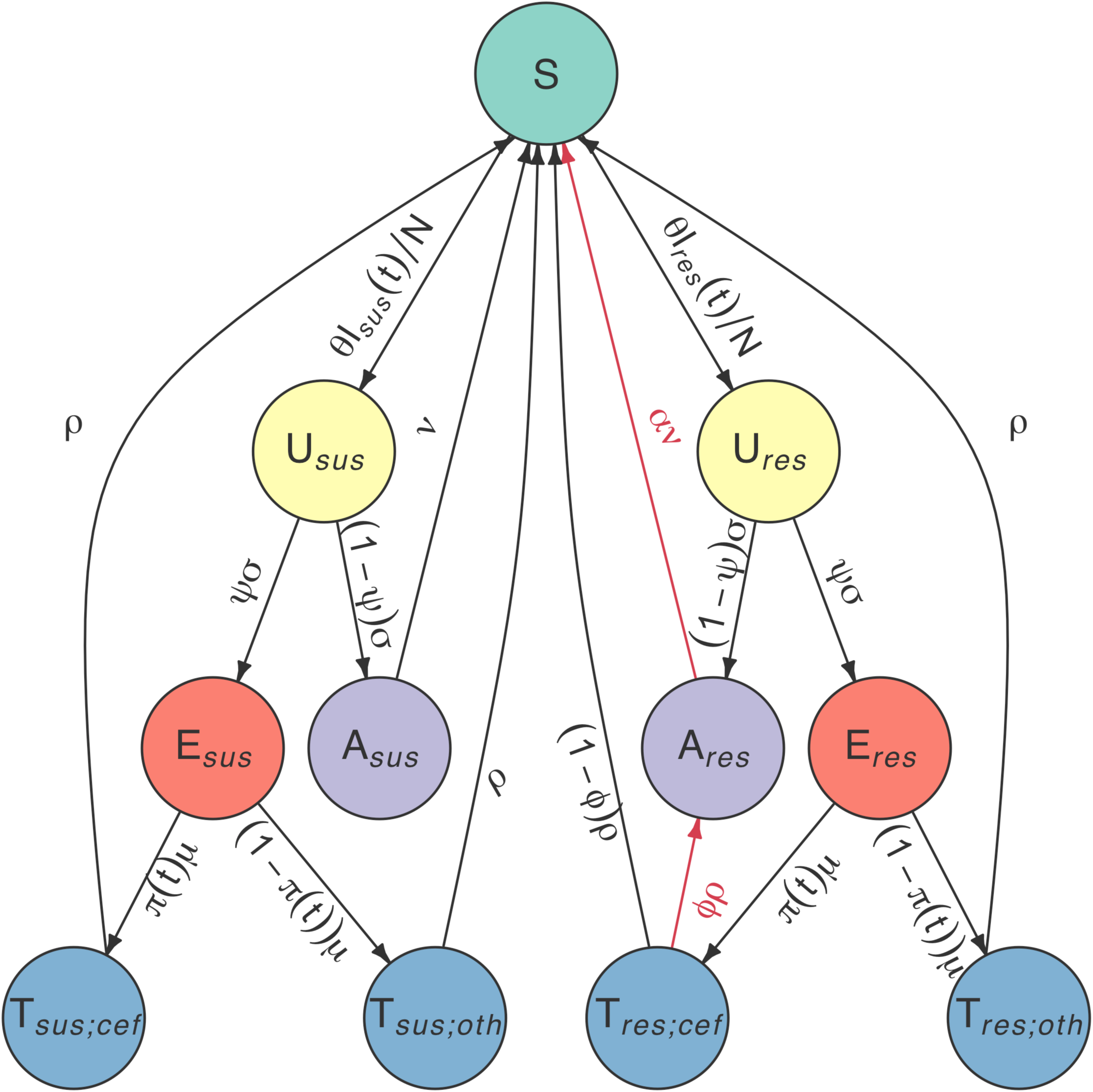
Flow diagram of model compartments with rates of transition between infection states. The left hand side represents infection with the cefixime-susceptible strains and the right hand side infection with the resistant strains. The two sides are symmetric with the exception of the two arrows highlighted in red which correspond to the cost and benefit of resistance.

**Table 1.** Parameter notations, prior and posterior distributions

| Parameter | Description | Prior | Posterior mean (95% CI) |
| --- | --- | --- | --- |
| $A_{\text{sus}}(0)$ | initial carriage of cefixime-susceptible infection | Uniform(0, ∞) | 630 (417, 888) |
| $A_{\text{res}}(0)$ | initial carriage of cefixime-resistant infection | Uniform(0, ∞) | 49 (19, 94) |
| $\theta$ | rate of transmission | Gamma(44, 10) | 5.2 (4.2, 6.5) |
| $\psi$ | proportion of infections that become symptomatic | Uniform(0, 1) | 62% (46%, 77%) |
| $\sigma$ | rate of departing incubation period | Gamma(17, 0.22) | 79 (46, 119) |
| $\nu$ | rate of recovery from asymptomatic infection | Gamma(8, 3.45) | 1.9 (0.9, 3.1) |
| $\mu$ | rate of seeking treatment when symptomatic | Gamma(3, 0.02) | 131 (33, 303) |
| $\rho$ | rate of recovery following treatment | Gamma(101, 1.9) | 53 (43, 63) |
| $\alpha$ | increased recovery from cefixime-resistant infection | Uniform(0, ∞) | 1.7 (1.4, 2.4) |
| $\phi$ | proportion of treatment failures for cefixime resistant infections treated with cefixime | Uniform(0, 1) | 84% (55%, 100%) |

### Calculation of the basic reproduction number

The basic reproduction number, *R*_0_, is a measure of the reproductive capacity of an infectious agent and is defined as the average number of secondary cases of gonorrhea arising from the introduction of a typical infected individual in a completely susceptible population. Where there is direct competition between strains, as in the situation we are modelling, the strain with the highest *R*_0_ will outcompete the others.

To calculate *R*_0_ we must consider the generation-time, defined as the expected time from an individual becoming infected to infecting another individual [37]. By considering the expected time spent in each compartment of the model corresponding to infection with the susceptible strain (ie. states *U*_sus_, *E*_sus_ and *A*_sus_ in Fig 2), we derive an analytical expression of the basic reproduction number 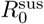 for the susceptible strains:

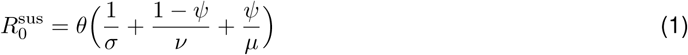

Similarly, the basic reproduction number 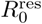 for the resistant strains is given by:

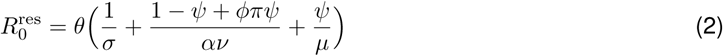

### Bayesian inference

We considered the data as a Partially Observed Markov Process, with the number of GUMCAD recorded cases, *Y*(*t*), and GRASP reported resistant cases, *Y*^res^(*t*), being the observed realisations of the underlying unobserved processes: the total incidence of gonorrhea infections, *Z*(*t*), and incidence of cefixime-resistant infections, *Z*^res^(*t*). The reporting process for the total number of cases recorded by GUMCAD was set to allow for a 10% under-reporting rate on average, with a 10% margin of error:

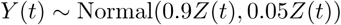
The probability of a cefixime-resistant case of gonorrhea being sampled by the GRASP study was assumed to be Poisson distributed with a sampling probability denoted *q*(*t*) derived from the coverage of the GRASP study over 2008 to 2014:

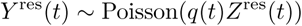
Based on these observations we aimed to infer the values of the ten parameters: *A*_sus_(0), *A*_res_(0), *θ*, *ψ*, *σ*, *v, α, μ*,*ρ* and *ϕ*.

An analytical expression for the likelihood of the observed data given our model is not available, so we obtained an unbiased estimate of the likelihood using a particle filter [38]. The estimated likelihood was then incorporated into a particle Monte Carlo Markov Chain (pMCMC) which was used to obtain a sample from the posterior distribution of the model parameters [39].

The model fitting was implemented using the R package pomp, which includes a pMCMC algorithm that can be used to perform Bayesian inference [40]. The algorithm was modified to enable parallel computation. The particle filter estimation of the likelihood was based on 1,000 particles, which was sufficiently robust to estimate the likelihood. 6 million iterations of the pMCMC were thinned by a factor of 100. Four separate chains were run with dispersed starting points, and compared using the R package coda [41]. The chains were found to have converged to the same posterior distribution based on the multivariate version of the Gelman-Rubin diagnostic, which was less than 1.1 for all inferred parameters [42, 43]. To ensure maximum robustness, the samples from the four chains were then combined, and found to have an effective sample size of more than 150 for all parameters.

### Prior distributions of parameters

Bayesian inference requires setting plausible priors for the model parameters. We used highly uninformative Uniform(0,1) priors for the two proportion parameters *ϕ* and *ψ* and Uniform(0,∞) priors for the three parameters *A*_sus_(0),*A*_res_(0)and *α*, which is an improper distribution but does not lead to an improper posterior distribution. For the five remaining parameters *θ*,*ν*,*σ*,*μ* and *ρ* we assigned informative Gamma priors based on literature review, as summarised in Table 1.

The transmission rate of infection, represented by the parameter *θ*, encompasses both the average number of sexual partners per year and the transmission probability per partnership. The Natsal-3 survey observed a mean number of sexual partners per year for MSM of 4.4 [34] and we would therefore expect *θ* to be slightly lower, to reflect the fact that not all contacts result in transmission. The prior distribution for *θ* was therefore set such that it was between 2.9 and 6.3 with 99% prior weight.

The expected duration of carriage for asymptomatic gonorrhea is not well measured. A study of 18 asymptomatic infected men saw no resolution in urethral infection in the 165 days until they received treatment [51]. Estimates of the duration of carriage in modelling studies have been based on calculations that take into account observed prevalence and assumed proportion of unobserved infection, and are often in the region of 6 months [52–54]. However, in recent work using genomic data, the greatest observed time to most recent common ancestor for bacterial genomes from known contact pairs was 8 months, suggesting that this estimate needs to be increased [30]. Duration of carriage may depend on infection site, for pharyngeal gonorrhea it has been estimated at 12 weeks, and for rectal infection has been estimated at one year [49, 55]. The parameter *v* was therefore assigned a prior that corresponded to a mean duration of carriage between three months and one year with 99% prior weight.

The duration of the incubation, symptomatic and treatment stages of infection have been estimated to be short, in the region of days rather than weeks [45–47]. Gamma priors were accordingly assigned to each of the three parameters *σ*,*μ* and *ρ*.

## Results

### Estimation of model parameters

We fitted our model of gonorrhea transmission to two different time series over the years 2008 to 2014: the total number of gonorrhea diagnoses in MSM in England [44] and the incidence of cefixime-resistant gonorrhea [14–20]. The posterior distribution of parameters shown in Fig 3 was obtained through Bayesian inference, implemented using a pMCMC method [39]. For each parameter we report the posterior mean estimate and 95% credibility interval shown in brackets (Table 1). The model suggests that at the end of 2007, the first year that cefixime-resistant cases were detected by GRASP [13], there were 630 cases (417, 888) of asymptomatic cefixime-susceptible gonorrhea (*A*_sus_(0)), and 49 cases (19, 94) of asymptomatic cefixime-resistant gonorrhea (*A*_res_(0)).

**Fig 3.**
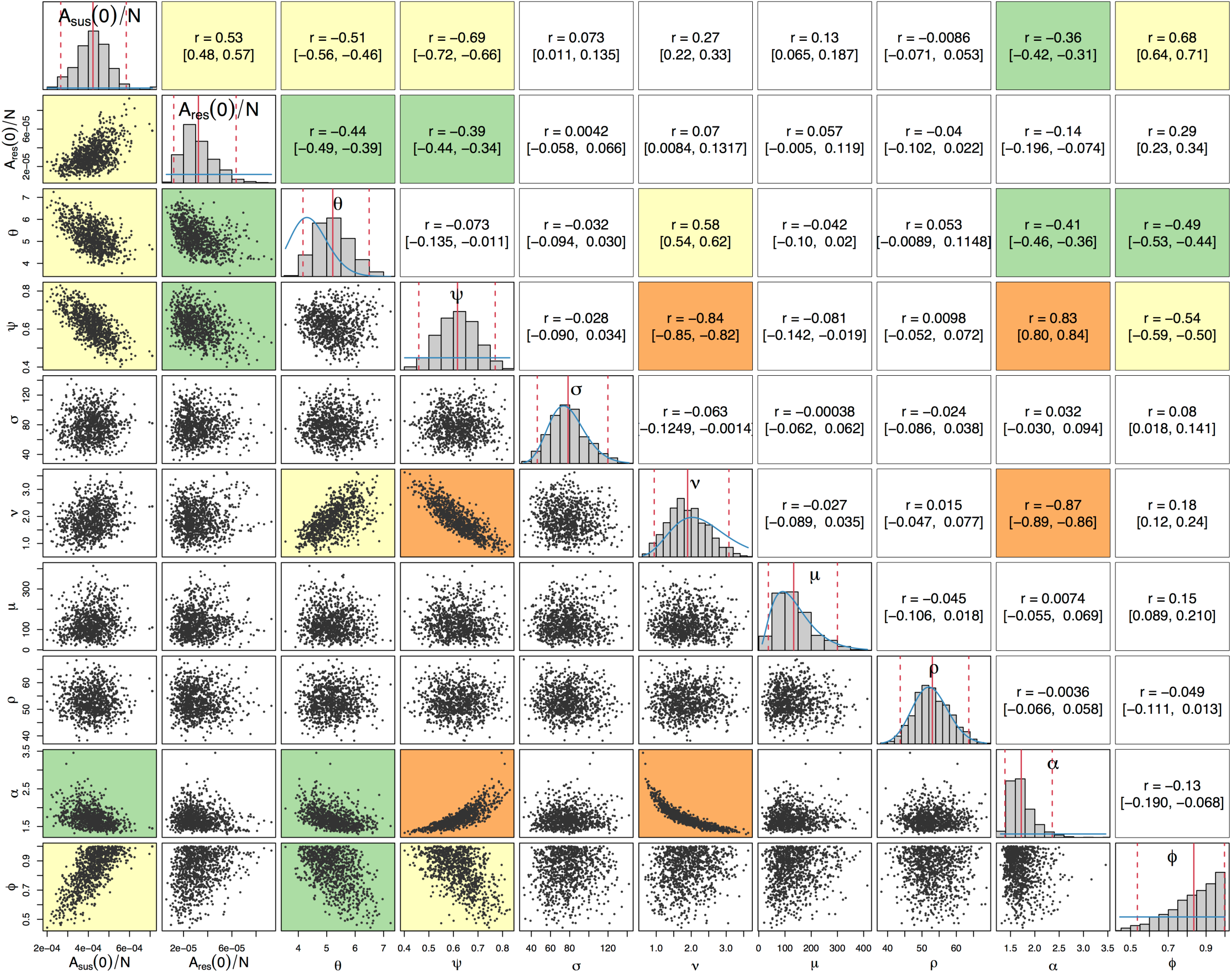
Posterior distributions of parameters. Diagonal plots show histograms of posterior distributions for all sampled parameters. Blue lines show prior distributions, red lines indicate posterior mean and 95% credible intervals. Plots below the diagonal show scatter plots based on 1,000 samples from the posterior, illustrating the relationships between pairs of estimated parameters. An orange background indicates a correlation higher than 0.8, a yellow background indicates a correlation between 0.5 and 0.8, a green background between 0.3 and 0.5, and a white background indicates a correlation less than 0.3. Plots above the diagonal show corresponding correlation coefficients with (95% CI).

The posterior distribution of the rate of transmission, *θ*, suggests a higher mean rate of infection than the prior expectation: 5.2 (4.2, 6.5), but the prior and posterior credible intervals overlap to a large extent, suggesting that the results are consistent with our prior knowledge. Our model predicts that the proportion *ψ* of infections that become symptomatic is 62% (46%, 77%). The three parameters *σ, μ* and *ρ*, corresponding respectively to the durations of the incubation period, symptomatic infection before seeking treatment, and the treatment phase, had posterior distributions similar to their prior distributions, indicating that the prior distributions were appropriate but that there is little additional information on these parameters in the data set. The posterior distribution of *v* has a slightly lower mean than prior,implying a longer mean duration of carriage of 193 days (118, 397). The prior and posterior credible intervals still intersect to a large extent so there is not significant evidence of a departure from the prior based on the data.

The last two parameters, *α* and *ϕ*, capture the difference between the susceptible and resistant strains in our model. The model predicts that in order to replicate observed incidence patterns recovery from asymptomatic cefixime-resistant gonorrhea occurs *α*=1.7 (1.4, 2.4) times faster than asymptomatic cefixime-susceptible gonorrhea, giving rise to a fitness cost. The model suggests a treatment failure proportion of (*ϕ*=84% (55%, 100%) for resistant gonorrhea treated with cefixime, so that resistance confers a fitness benefit in an environment in which cefixime is highly prescribed.

Beyond the marginal posterior distributions of the parameters described above, it is informative to study their posterior correlations. The pairwise posterior relationships between the ten parameters are depicted in Fig 3. Parameters *σ, μ* and *ρ* did not show a strong correlation with any parameters; as expected the short duration of the incubation, symptomatic and treatment stages of infection led to these parameters contributing relatively little to the dynamics of infection. The strongest correlation was found between *ν* and *α*, -0.87 (-0.89, -0.86), corresponding to the trade-off required to maintain the duration of carriage of resistant infection, which is equal to 1/(*αν*). Parameters *v* and *ψ* were also highly negatively correlated, -0.84 (-0.85, -0.82), which corresponds to the trade-off between duration of carriage, 1/*ν*, and the proportion of infections entering the carriage state, (1 – *ψ*). The two negative correlations of both *α* and *ψ* with *ν* lead to a positive correlation between *α* and *ψ*.

### Posterior predictive analysis

The total number of gonorrhea cases in MSM observed by GUMCAD and the number of cefixime-resistant infections isolated in MSM by GRASP were compared with simulated datasets using parameters sampled from their posterior distributions. Fig 4 demonstrates that the simulated data closely emulate the real data. The real data are within the 95% predictive intervals at all time points, indicating a good fit of the model to the data [57].

**Fig 4.**
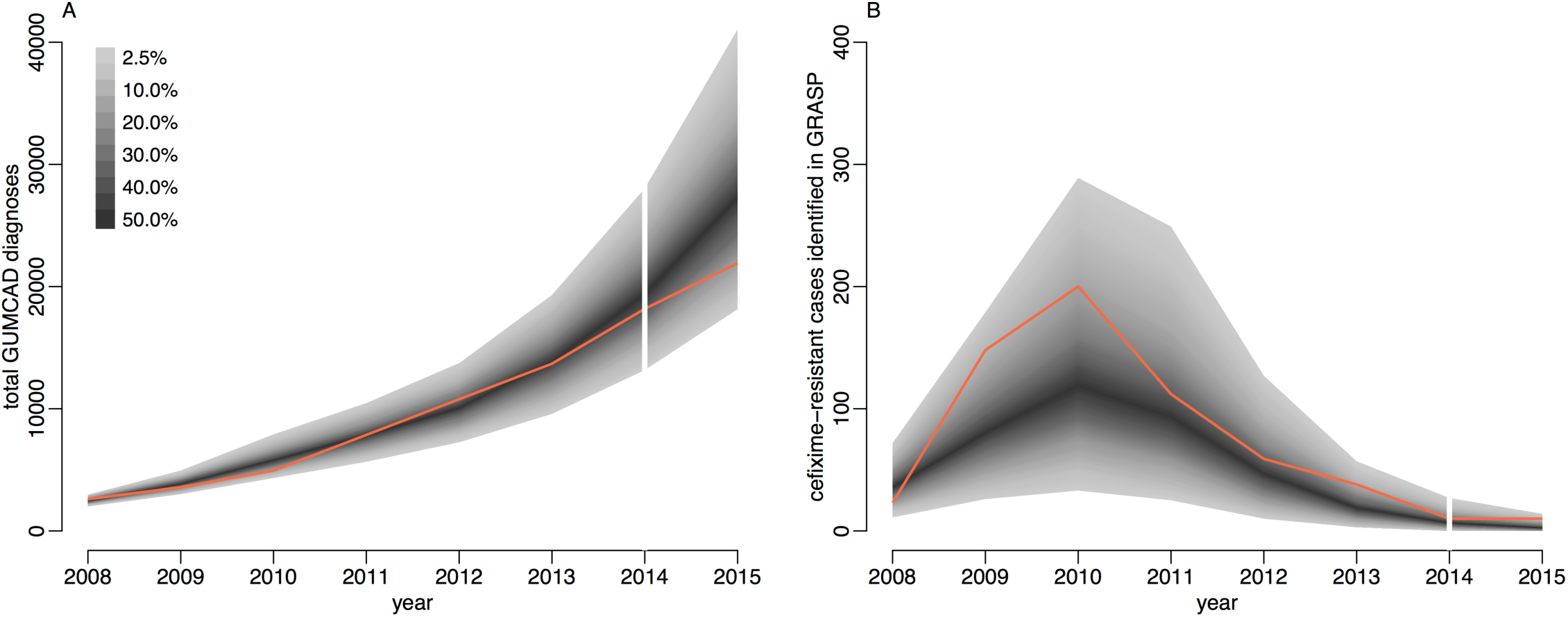
Comparison of simulated and observed cases of gonorrhea. Panel A shows total number of cases and panel B shows only the cefixime-resistant cases. Observed data are shown in orange, with the shaded area showing the 95% posterior predictive interval (based on 1,000 simulations using samples from posterior distribution).

The total number of cases of gonorrhea observed by GUMCAD, and the number of cefixime-resistant cases observed by GRASP in 2015 [21] were not used in the model fitting process, and were used to provide an independent check of the model fit. Both data points are within the 95% probability intervals predicted by our model: 21,915 gonorrhea diagnoses were recorded by GUMCAD, compared to 27,668 (18,144, 41,054) diagnoses predicted by the model; 10 cefixime-resistant cases were recorded by GRASP, compared to 4 (0, 14) cases predicted by the model. Our modelling suggests that in 2015 0.7% (0.4%, 1.1%) of MSM in England may be carriers of asymptomatic gonorrhea.

### Comparative analysis of basic reproduction numbers

A key threshold in epidemic theory associates the persistence of disease in a population with a basic reproduction number greater than one [58]. Using Eq 1 we obtain a posterior estimate for the basic reproduction number for cefixime-susceptible infection of 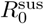 which suggests that the cefixime-susceptible strain of gonorrhea is expected to persist in the population without further intervention (Fig 5A). Under our hypothesis the basic reproduction number for cefixime-resistant gonorrhea depends on the frequency of cefixime prescription (Eq 2) and so can be considered as a function 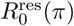 where (*π*) is the proportion of gonorrhea diagnoses being treated by cefixime. In the two extreme cases when no cefixime is prescribed (*π* = 0, meaning that treatment is always effective) and only cefixime is prescribed (*π* = 1, meaning that only a proportion 1 – *ϕ* of treatment is effective) we estimate a basic reproduction number for resistant gonorrhea of 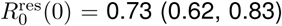 and 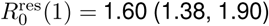, respectively. At its height in 2008 the frequency of cefixime prescriptions was 70%, we estimate that at this time the basic reproduction number for resistant gonorrhea was 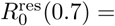 1.34 (1.22, 1.51). The former estimate is ¡ 1 whereas the latter two are ¿ 1, which is consistent with the fact that between 2005 and 2010, when cefixime was often used to treat gonorrhea, resistance to cefixime increased whereas with the discontinuation of cefixime usage from 2011 resistance has decreased.

**Fig 5.**
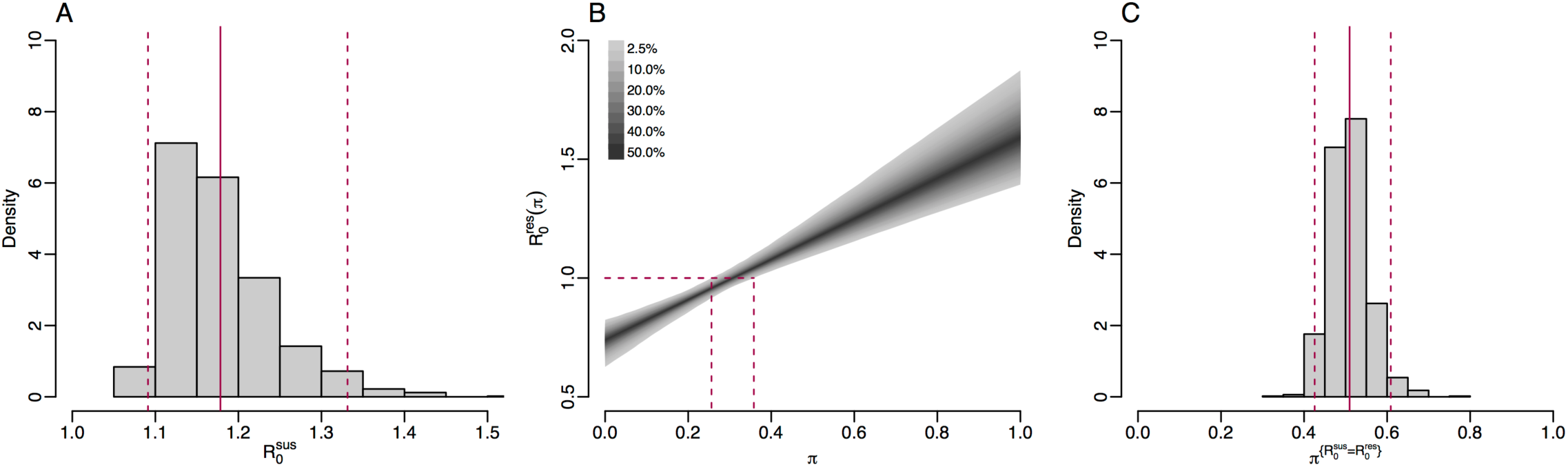
**A**. histogram of posterior estimate of 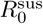 **B**. 95% credible interval of 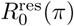 against *π* with dashed lines showing 95% credible interval for 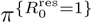 **C**. histogram of posterior estimate of 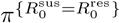: the threshold of cefixime prescriptions above which 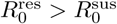

We can estimate the frequency of cefixime prescriptions above which we expect the resistant strain to persist, corresponding to when the fitness benefit of cefixime resistance is greater that its fitness cost, by setting 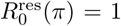 and solving for *π* in Eq 2. We denote this threshold 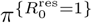, and thus obtain a posterior estimate of 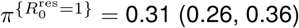 (Fig 5B). This result suggests that up to a quarter of gonorrhea treatments could be with cefixime monotherapy without causing a cefixime-resistant epidemic. Another important threshold is the level of cefixime prescriptions above which the resistant strain of gonorrhea is fitter than the susceptible strain. We denote this threshold 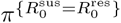.By setting 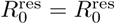 and equating Eqs 1 and 2 we obtain a posterior estimate of 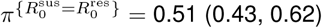 (Fig 5C).

### Impact of cefixime usage on simulated resistance trends

The basic reproduction numbers derived above are informative, but do not capture completely the complex dynamics of infection transmission that occur when accounting for stochasticity, competition between susceptible and resistant strains, and non-negligible fractions of the population becoming infected. To further study the impact of cefixime prescribing on the cefixime-resistant and susceptible epidemics, we performed stochastic model simulations over eight years from 2008 to 2015 using parameters drawn from their posterior distributions and examining scenarios with a constant frequency of cefixime prescriptions ranging from no use of cefixime (*π* = 0) to all gonorrhea cases being treated by cefixime (*π* = 1). Fig 6A shows that, when more than 35% of gonorrhea cases were treated with cefixime, there was a 95% probability that the resistant outbreak persisted in 2015. This is comparable to our estimate of 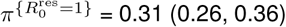 (Fig 5B), the level of prescriptions above which the fitness 232 benefit of cefixime resistance is greater than the fitness cost.

**Fig 6.**
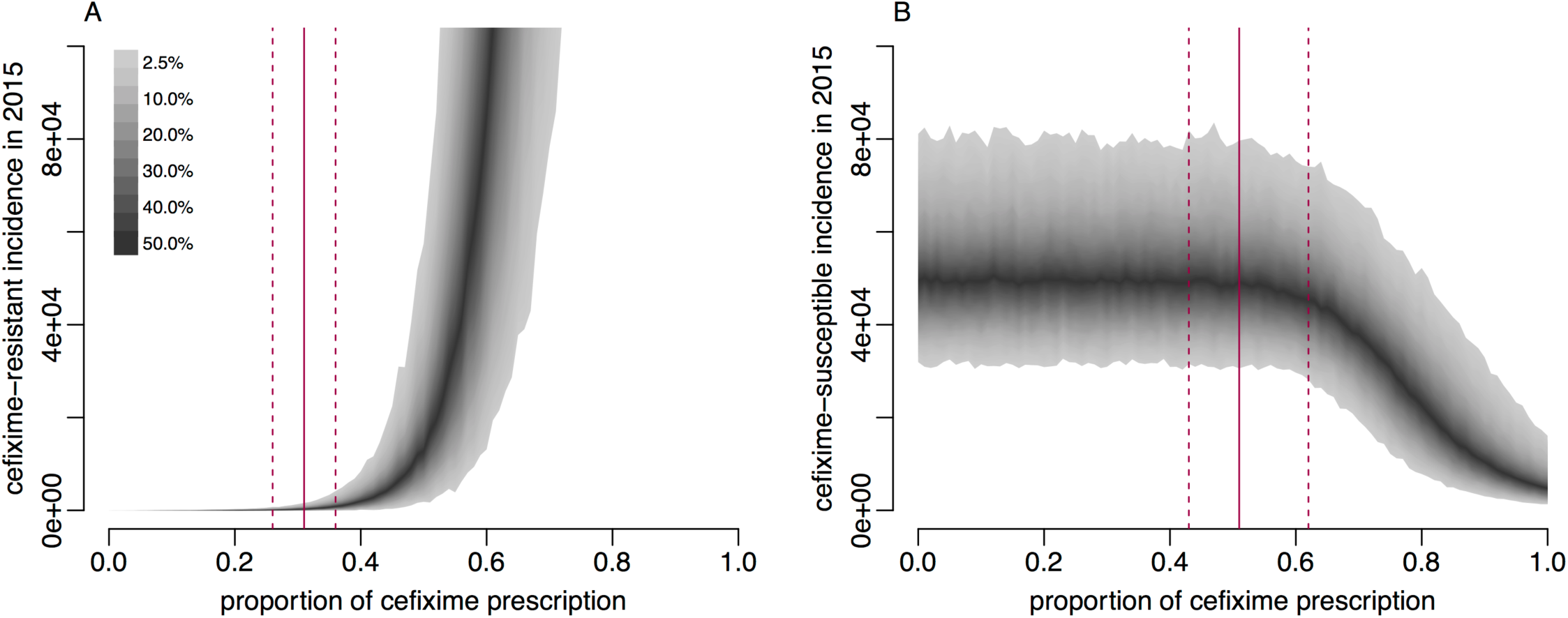
Incidence of gonorrhea in 2015 based on simulations from 2004 to 2015 with varying levels of cefixime prescribing. **A**. incidence of the cefixime-resistant strain, red lines show 95% credible interval for 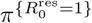. B. incidence of the cefixime-susceptible strain, red lines show 95% credible interval for 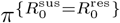 Shaded areas show the 95% posterior predictive intervals (based on 1,000 simulations using samples from posterior distribution).

Fig 6B shows that, when more than 50% of gonorrhea cases were treated with cefixime, the simulated incidence of cefixime-susceptible infection began to fall, with the cefixime-resistant strain becoming more common. This supports our analytical estimate of 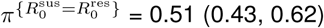 (Fig 5C), the level of cefixime prescriptions above which the resistant strain becomes fitter than the susceptible strain.If cefixime were used to treat more than 60% of cases then the level of cefixime resistance would become greater than 50% at the end of the eight year simulation period and if cefixime were used to treat all cases resistance would be close to 100%.

## Discussion

We have used mathematical modelling and Bayesian inference methods to uncover insights into the dynamics of cefixime resistance in gonorrhea. We quantified both the fitness cost and fitness benefit of resistant strains, which allowed us to make predictions about the future prevalence of resistance as a function of how often cefixime is prescribed. Our results indicate that cefixime could be used to treat uncomplicated cases of gonorrhea without incurring the risk of causing a resistant epidemic like the one that happened in 2007-2012, provided its frequency of use were controlled, enabling continued use of an ‘abandoned’ antibiotic. Our analysis suggests that cefixime could be used to treat up to 25% of cases, but this threshold should be used cautiously for reasons described below.

Our modelling approach requires making a number of assumptions, and it is important to consider their suitability. For example, it was assumed that all cefixime-susceptible infections were cured, regardless of which antibiotic was prescribed. The prescription data shows that between 2008 and 2015 the vast majority of non-cefixime prescriptions were for ceftriaxone, either alone or in combination with azithromycin, so the assumption of cure is reasonable given that ceftriaxone resistance reports remain sporadic in England. Our model implicitly assumes that there is no co-infection with both strains,and no evolution of resistance happening within-host. This simplification has been used in a number other studies on the epidemiology of antimicrobial resistance [61, 62]. Ignoring within-host competition between resistant and susceptible strains following co-infection is justified here by the fact that both strains have low prevalence, making co-infection very unlikely. Within-host evolution of resistance was included in a recent gonorrhea modelling study [67] but clearly this is a rare event which only increases by one the number of resistant infections, which we estimated to be already equal to 49 at the start of the epidemic simulation on the 1st January 2008.

Our model also makes assumptions concerning the cost and benefit of cefixime resistance. The fitness cost of the mutation conferring resistance is assumed to be constant over time, however compensatory mutations have been observed in other bacterial pathogens that reduce the initially high fitness cost of antibiotic resistance [59, 63]. It is clear from our analysis that there was a substantial fitness cost to cefixime resistance when the prescription protocol was changed in 2010, which is the reason why the resistance level subsequently fell. We cannot rule out that compensatory mutations took place after resistance initially emerged, but this would mean that the initial cost was even higher and in these conditions resistance would have been unlikely to emerge at all. Our formulation of the dynamics of the 13 fitness cost of resistance was via a reduction in the duration of asymptomatic carriage. In the absence of evidence of the resistance mechanism the fitness cost could plausibly be modelled through reduced transmissibility of the resistant strain [67], which would not affect our overall conclusions, in particular regarding the basic reproduction number analysis and predictions of the impact of cefixime usage on future resistance trends.

The ceftriaxone-azithromycin dual therapy is currently effective, but it represents a last resort, so that we urgently need a strategy for what would be done if it stopped working. It is likely to be just a matter of time before this happens, with the first reported failure of the dual therapy having occurred in 2015 [65].Resistance to azithromycin was detected in a recent outbreak that started in the North of England [5], and is now reported in almost 10% of tested isolates [21]. Resistance to ceftriaxone remains rare, but minimum inhibitory concentration (MIC) levels have been steadily increasing [66]. If alternative treatment options could be used, even for a minority of cases, then it would delay and maybe even prevent the emergence of resistance to the dual-therapy antibiotics by reducing the fitness benefit it would confer.

For some previously used antibiotics, such as penicillin or ciprofloxacin, significant levels of resistance remain in the gonococcal population (24% and 39% in 2015, respectively [21]), so that they could not be recommended even for a small fraction of cases. These antibiotics could be prescribed only if drug-sensitivity could be quickly established, for example using real-time PCR assays [70, 71], or whole genome sequencing [64, 72], which both remain experimental. In contrast, the fact that resistance to cefixime has become very low in England (around 1% in 2015, [21]) makes it a prime candidate for return into action without the need for case-by-case susceptibility testing. Perhaps the greatest threat posed by this proposed strategy would be the evolution of compensatory mutations which could reduce the fitness cost of resistance. As previously mentioned, compensatory mutations do not seem to have emerged during the 2007-2012 cefixime-resistant epidemic, but if they did the acceptable prescribing proportion would be lowered, and the probability of persistence of cefixime resistance increased. Therefore, a redeployment of cefixime would require close monitoring of resistance trends in England [68] and beyond [73, 74].

## Supporting information

### S1 Appendix

**Model equations and stochastic simulations.**We developed a simple susceptible-infected-susceptible (SIS)-type stochastic compartmental model that describes the transmission and natural history of gonorrhea, as illustrated in Fig 1 with parameters summarised in Table 1. The population was assumed to be closed due to the short time period under consideration. A deterministic version of the model is described by the following differential equations.

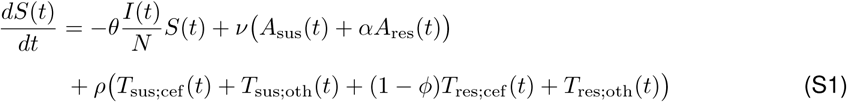

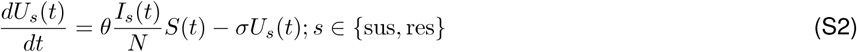

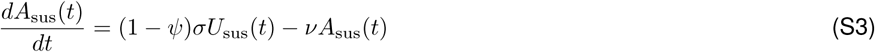

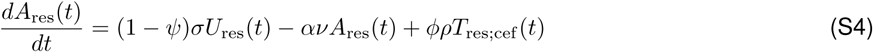

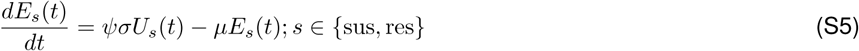

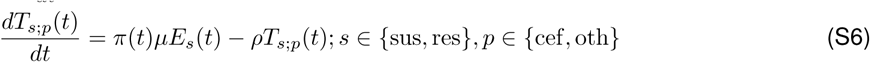

We used a stochastic version of the model described in Eqs S1-S6. The process was initialised on 31 December 2007, with an initial number of asymptomatic infections *A*_s_(0) for *s* ∈ {sus, res}, and the remainder of the population being susceptible. Simulation proceeds through repeated iterations of the steps below for each day. Transition variables *d*_1_, …,*d*_17_ are drawn from the following distributions:

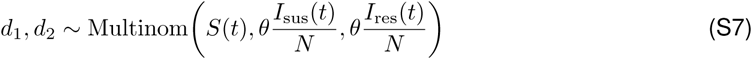

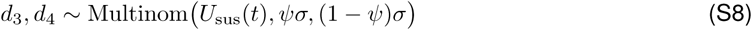

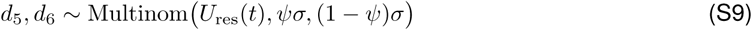

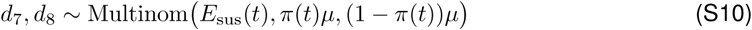

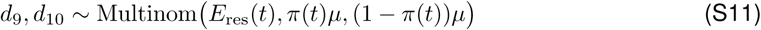

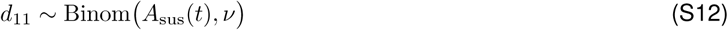

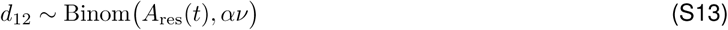

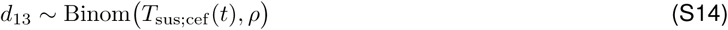

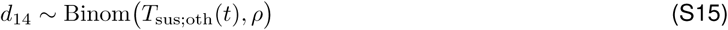

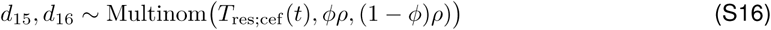

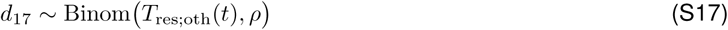

The compartments of the model are then updated as follows:

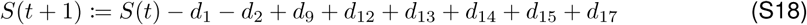

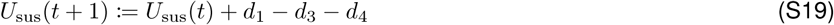

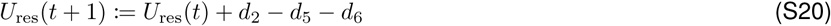

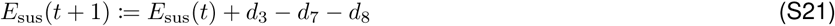

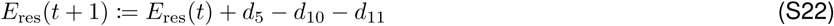

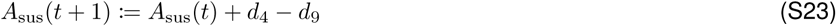

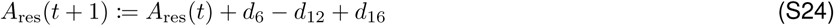

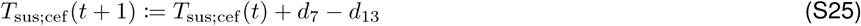

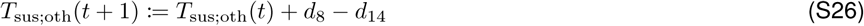

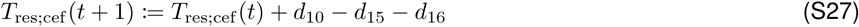

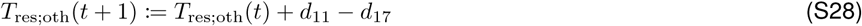

## Acknowledgments

We wish to thank David Eyre, John Paul and Brian Spratt for stimulating discussions and constructive comments on an earlier version of this manuscript.

